# Tailoring CRISPR-Cas Immunity for the Degradation of Antibiotic Resistance Genes

**DOI:** 10.1101/2022.03.09.483686

**Authors:** Xin Li, Nan Bao, Zhen Yan, Xian-Zheng Yuan, Shu-Guang Wang, Peng-Fei Xia

**Affiliations:** Shandong Key Laboratory of Water Pollution Control and Resource Reuse, School of Environmental Science and Engineering, Shandong University, Qingdao 266237, China; Sino-French Research Institute for Ecology and Environment (ISFREE), Shandong University, Qingdao 266237, China

**Keywords:** CRISPR, antibiotic resistance genes, conjugation, synthetic biology, environmental biotechnology

## Abstract

The evolution and dissemination of antibiotic resistance genes (ARGs) are prompting severe health and environmental issues. While environmental processes are key barriers preventing the spread of ARGs, they are often sources of ARGs at the same time, as ARGs may be required and accumulate in the biological treatment units. An upgrading of environmental biotechnology is imperative and urgent. ARGs confer antibiotic resistance based on the DNA sequences rather than the chemistry of DNA molecules. An ARG can be considered degraded if its sequence was disrupted. Therefore, we present here that CRISPR-Cas immunity, an archaeal and bacterial immune system for eliminating invading foreign DNAs, can be repurposed and tailored for the degradation of ARGs. By deploying an artificial IncP machinery, the designed system, namely VADER, can be successfully delivered via bacterial conjugation. Then, we propose a new sector for ARG degradation to be implemented as a complement to the biological units in the framework of environmental processes. In this endeavor, a prototype conjugation reactor at a 10-mL-scale was devised, and 100% of the target ARG were eliminated in the transconjugated microbes receiving VADER in the reactor. By generating a nexus of synthetic biology and environmental biotechnology, we believe that our work is not only an enterprise for tackling ARG problems but also a potential solution for managing undesired genetic materials in general in the future.

**Importance:** Antibiotic resistance has been causing severe health problems and leading to millions of deaths in recent years. Environmental processes, especially the wastewater treatment sector, are important to barrier the spread of antibiotic resistance from the pharmaceutical industry, hospitals, or civil sewage. However, they have been identified as the source of antibiotic resistance at the same time, as antibiotic resistance with its main cause antibiotic resistance genes (ARGs) may be required and accumulate in the biological treatment units, leading to the dissemination of ARGs. Here, we transplanted the CRISPR-Cas system, an immune system via programmable DNA cleavage, to environmental biotechnology for tackling the antibiotic resistance dilemma thereof, and we propose a new sector in environmental processes specialized in ARG removal with a reactor inhabiting the CRISPR-Cas system per se. Our study provides a new angle to resolve public health issues via the implementation of synthetic biology at the process level.

## Introduction

Antibiotic resistance is a threatening issue for human health (1,2). Approximately, 4.95 million deaths were associated with antibiotic resistance in 2019, while 1.27 million deaths were direct consequences of antibiotic resistance (2). Most antibiotic resistance results from antibiotic resistance genes (ARGs), which encode machinery that inactivate or pump out antibiotics. An ARG-harboring pathogen is more harmful to humans as the correlated antibiotic is no longer effective. In the environment, ARGs can disseminate via horizontal gene transfer or evolve under the selection of antibiotics, leading to increased health and environmental concerns (1). Though many efforts have been made to understand and resolve ARG-related issues, fine control of ARGs has not yet been implemented.

Environmental processes, i.e., wastewater treatment, are key barriers that prevent ARGs from entering the environment (3). It combats antibiotic resistance by degrading antibiotics, killing antibiotic resistance microbes, and directly removing ARGs. However, detection of various types of ARGs has been reported in and near wastewater treatment facilities (4–8), indicating them potential sources for ARGs at the same time. It, unfortunately, seems inevitable as the ARGs may be required and accumulate for the removal of antibiotics in the wastewater from pharmaceutical industries, hospitals, or even municipal sewerage systems, leading to a dilemma in addressing ARG problems. To tackle this challenge, environmental scientists have interrogated the fate of ARGs in engineered and natural systems (9–12), and have been developing remediation methods (13–15). For current environmental applications, most of the methods remove ARGs as biomolecules based on the physiochemistry of DNAs (13). Yet, a fragment of DNA molecule is considered an ARG due to the sequence, which encodes proteins that confer resistance to the corresponding antibiotics. Thus, an ARG can be considered degraded when the sequence was disrupted.

CRISPR-Cas system is an archaeal and bacterial immune system that has been repurposed as gene-editing tools (16–19). As an immune system, the CRISPR-Cas system finds and cleaves a target DNA under the lead of a guide RNA (gRNA). As such, by applying the programmable CRISPR-Cas immunity, ARGs can be deliberately degraded. Pioneer studies have validated that the CRISPR-Cas system can eliminate ARGs for medical applications (20–23), and the system can be delivered via transduction or conjugation (22, 23). We envision that the CRISPR-Cas system can be adapted and implemented in environmental bioprocesses for ARG removal. It is possible to instrument a controllable new sector (e.g., a conjugation reactor) harnessing the CRISPR-Cas immunity programmed for the degradation of ARGs. The new sector may serve as a complement to the biological processes, where ARGs in the microbes going through this unit can be eliminated in vivo. Then, the “clean” microbes may return to their assigned bioprocess and will be safer for disposal when required.

Thus, we devised and demonstrated VADER, an en<u>v</u>ironmentally-<u>a</u>imed degradation system for <u>e</u>liminating A<u>R</u>Gs, based on the *Streptococcus pyogenes* CRISPR-Cas system and the IncP conjugation machinery. We designed VADER to work in a one-plasmid and constitutive mode to avoid the complexity of multi-plasmids and extra induction steps. Then, a streamlined method was optimized for generating VADER plasmids. Finally, we devised a prototype conjugation reactor and observed the successful delivery of VADER and the degradation of ARGs in the reactor. Our study shows a novel path to tackle ARG problems in environmentally relevant contexts with CRISPR-Cas immunity at the process level.

## Results

### Modular design and demonstration of VADER

*S. pyogenes* CRISPR-Cas immunity works in an RNA-guided manner and cleaves the target DNA with the single effector Cas9 and a programmed gRNA **(Figure 1A)** (24). To employ this machinery, we designed VADER as a one-plasmid system with constitutive functions for simplicity and efficiency to avoid incompatibility and complexity of multi-plasmids or extra induction steps. First, we constructed a plasmid pCasEnv with *cas9* from *S. pyogenes* as the ARG degradation module, a selection marker *(gentR),* and a broad-host *pBBR* origin of replication (ori) to expand the application spectrum (25). Meanwhile, gRNA01 was generated via inverse PCR with pMV-gRNA as a template **(Table S1 and Table S3)**. The gRNA01 cassette consisting of the J23119 promoter, a spacer and the chimeric scaffold was fused to pCasEnv, generating pEL01 as the first VADER plasmid **(Figure 1B)**. Then the following plasmids were generated in a streamlined method by inverse PCR with pEL01 as a template and programmable primer pairs to replace the spacer **(Figure 1B)**.

**Figure 1.**
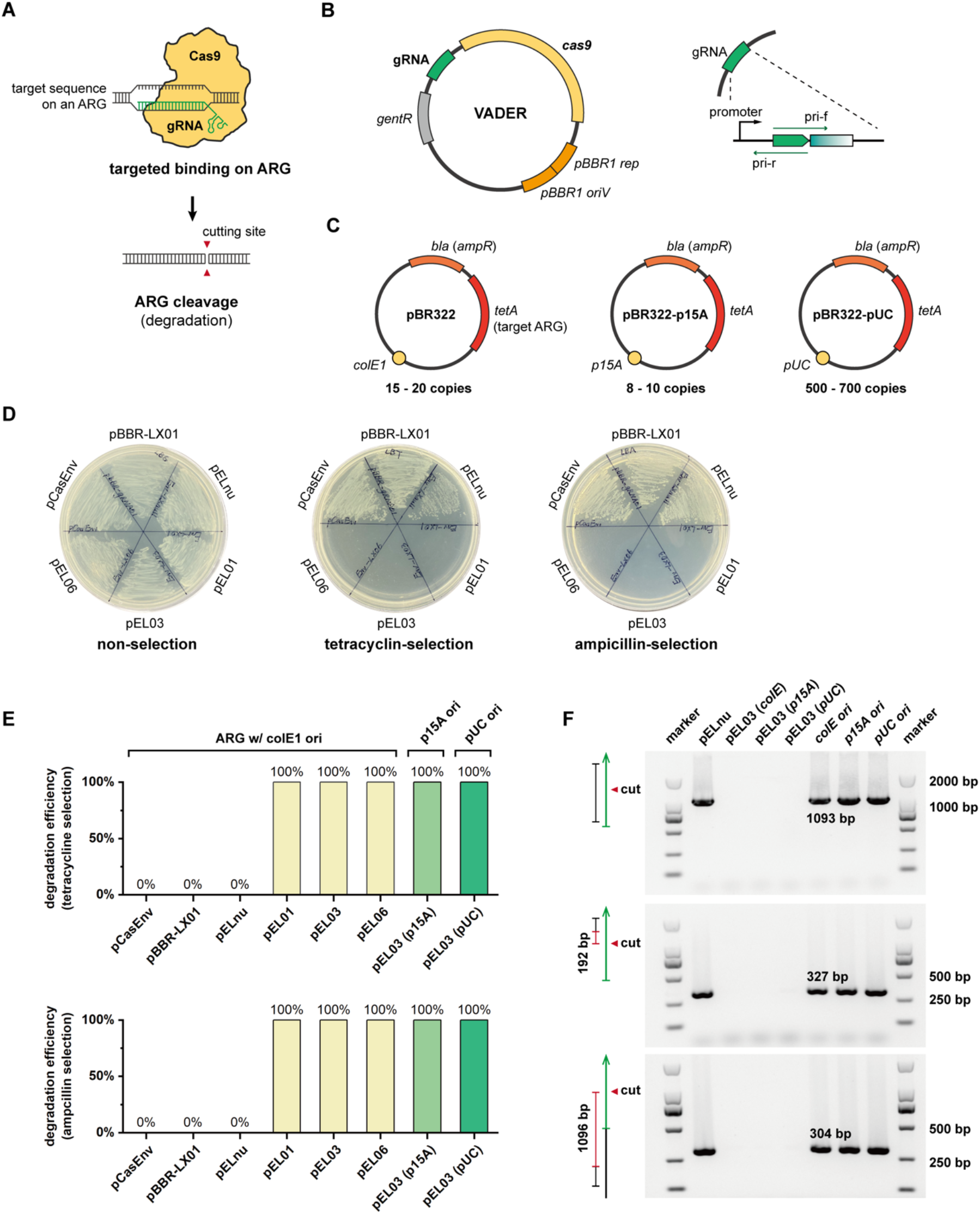
Design and demonstration of VADAR. (A) Working mechanism of CRISPR-Cas immunity on ARG degradation. (B) Design scheme of VADER plasmids and the streamlined generation of VADAR plasmids. The back-to-back primer pairs shown as pri-f and pri-r are used for quick exchange to spacer sequences. (C) Maps of target plasmids harboring the target ARG *tetA* with different copy numbers. (D) Demonstration of VADER plasmids pEL01, pEL03, and pEL06 for *tetA* degradation and the validation of plasmid integrity with pCasEnv, pBBR-LX01, and pELnu as controls. The transformants were first selected on plates with gentamycin to ensure the transformation of VADER plasmids. Then, colonies were randomly picked and streaked on plates with tetracycline and ampicillin. (E) Degradation efficiencies of VADER were calculated via evaluation of tetracycline resistance and ampicillin resistance of randomly picked colonies from the transformants with VADER plasmids. (F) Electrophoresis documentation of PCR fragments with primer pairs designed for amplification of the intact *tetA*, small fragments nearby (192 bp of distance) and away from (1096 bp of distance) the cleavage site by VADER.

As most of the ARGs are plasmid-borne (1,26), we focused on and evaluated VADER for eliminating ARGs on plasmids. We chose pBR322 as the target plasmid with a *tetA* gene conferring tetracycline resistance (Tet^R^) and a *bla* gene conferring ampicillin resistance (Amp^R^) **(Figure 1C)**. *tetA* was regarded as the target ARG, while *bla* was used to check the integrity of the target plasmid. Three gRNAs, gRNA01, gRNA03, and gRNA06, were generated with spacers targeting different sites on the coding or non-coding strand of *tetA* **(Table S4)**. We found that, by applying VADER, the *tetA* gene was efficiently inactivated. All resulting transformants receiving VADER plasmids pEL01 (gRNA01), pEL03 (gRNA03), and pEL06 (gRNA06) **(Table S1 and Table S3)** lost Tet^R^ and Amp^R^ **(Figure 1D and 1E)**, indicating the degradation of *tetA* and the elimination of pBR322. Under the current design, a VADER plasmid contains *pBBR* ori which shows 8 – 10 copies of plasmids in a cell, and the target plasmid pBR322 harbors *colE1* ori, which gives 15 – 20 copies of *tetA* (27). To further explore the capability of VADER, we designed two more plasmids with varied copy numbers by replacing *colE1* ori with *p15a* ori and *pUC* ori, generating pBR322-p15a and pBR322-pUC, respectively. pBR322-p15a presents 8 – 10 copies of *tetA* per cell, and pBR322-pUC gives 500 – 700 copies of *tetA* per cell (28). We transformed pEL03 to *E. coli* with pBR322-p15a and pBR322-pUC and observed successful elimination of *tetA* on these plasmids **(Figure 1E)**. Normally, mobilizable plasmids, such as RP4-2 and RK2, are low-copy plasmids with less than 10 copies in a cell (26, 29). As VADER managed to degrade ARG carried by a plasmid with a copy number up to 700, it shows sufficient capability for degradation of plasmid-borne ARGs.

Furthermore, we tested whether *tetA* was actually degraded. Three pairs of primers were designed to detect **1)***tetA* as an intact gene with a 1093-bp fragment, **2)**a small fragment (327 bp) near the cleavage site, and **3)**another small fragment (304 bp) that has a distance of 1095 bp from the cleavage site **(Figure 1F)**. We transformed non-targeting (pELnu) and targeting (pEL03) VADER plasmids to *E. coli* with either pBR322, pBR322-p15a, or pBR322-pUC, and randomly picked the transformants for PCR analysis. We identified the PCR fragments with all three primer pairs for the transformants with the non-targeting VADER and for the positive controls (purified plasmids), but no PCR fragments for the colonies receiving the targeting VADER could be detected **(Figure 1F)**. These results demonstrated that *tetA* was degraded via a single cleavage in the middle, and the resulting degradation covered the whole length of *tetA*.

### Customizing and duplexing of gRNAs

Despite the demonstration of VADER, we encountered difficulties in generating VADER plasmids via the streamlined inverse PCR method. In some cases, the number of transformants was quite limited, and the plasmids from these transformants often contained mutations. These might result from the toxicity of foreign CRISPR-Cas systems in bacteria (30, 31), and the toxicity as a selective pressure drove mutations in the CRISPR-Cas system. Previous reports highlighted the toxicity of CRISPR-Cas in bacteria (31–33), and, specifically, Beisel and colleagues noticed a high mutation rate in generating plasmids for CRISPR-Cas-based base editing (31). For better-streamlined plasmid generation, further optimization of VADER is necessary.

We examined the mutation patterns in the mutated plasmids thereof and found most of the mutations lie in the TATA box of the J23119 promoter leading gRNAs in our design **(Figure 2A)**. We tested a representative mutated plasmid when generating pEL14 (gRNA14, **Table S3**), with TAAAT instead of TATAAT as the TATA box and surprisingly observed high efficiency of ARG degradation as well **(Figure S1)**. We hypothesized that the mutated J23119 promoters have lower strength and alleviate the toxicity of CRISPR-Cas systems, making them better candidate promoters. Two new promoters J23119* and J23119** were designed with one nucleotide deleted at the position −12 and −11, respectively. The pEL03 plasmid was re-constructed at ease with these new promoters **(Figure 2B)**, and both constructs showed the same efficiency (100%) in *tetA* degradation compared to the original J23119 promoter **(Figure 2C)**. We demonstrated the customized J23119** promoter with two more VADER plasmids, pEL12 and pEL13, both giving 100% efficiency of *tetA* degradation **(Figure S2)**. Hereafter, we employed the J23119** promoter for the generation of VADER plasmids, and our results also supported that promoter engineering for fine-tuning gRNA levels will be a solution for alleviating CRISPR-Cas toxicity in bacteria.

**Figure 2.**
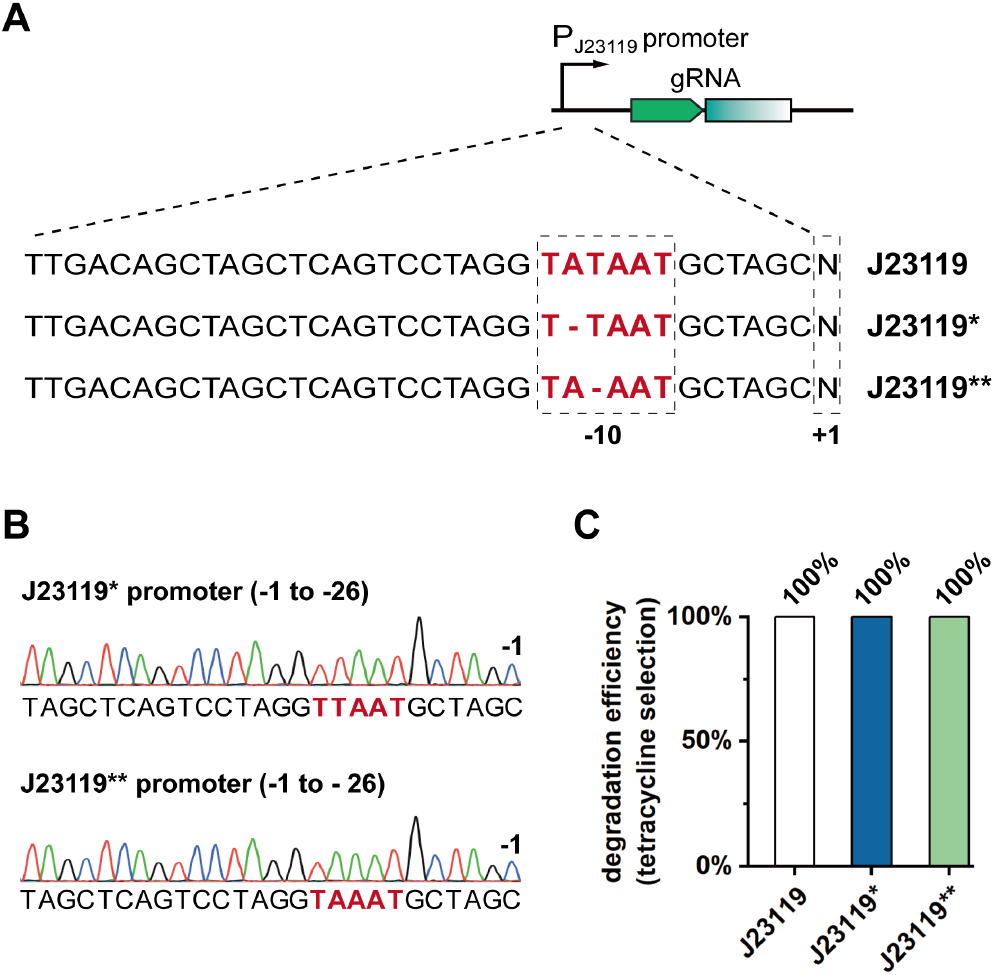
(A) Mutation patterns in J23119 promoter of mutated VADER plasmids. (B) Validation of re-generated pEL03 with J23119* and J23119** promoters. TATA box regions were highlighted in red. (C) Degradation efficiency of VADER plasmid pEL03 with J23119, J23119* and J23119** promoter driving gRNA03, respectively.

Moreover, we observed the low on-target efficiency of VADER for certain target sequences. A dual gRNA system is necessary for efficient degradation of ARGs in case one of the gRNAs is not functional. As an example, we generated a VADER plasmid pELdu with tandem gRNAs, gRNA11, and gRNA03 **(Figure 3A)**. In a preliminary test, we found that pEL11 with gRNA11 cannot degrade *tetA* **(Figure 3B)**. This may be a result of DNA methylation or protospacer specificity. When we employed pEL11 for *tetA* degradation in methylation deficient *E. coli* (*dam*^-^ and *dcm*^-^), we found that *tetA* was efficiently eliminated in half of the transformants **(Figure 3B)**, implying a combined mechanism underlying the low on-target efficiency for certain protospacers. Then, we applied pELdu in *E. coli* with functional methylation mechanism and methylation deficiency. One hundred percent efficiency of *tetA* degradation was observed in all tests, indicating that our duplex system worked well as double insurance for ARG degradation.

**Figure 3.**
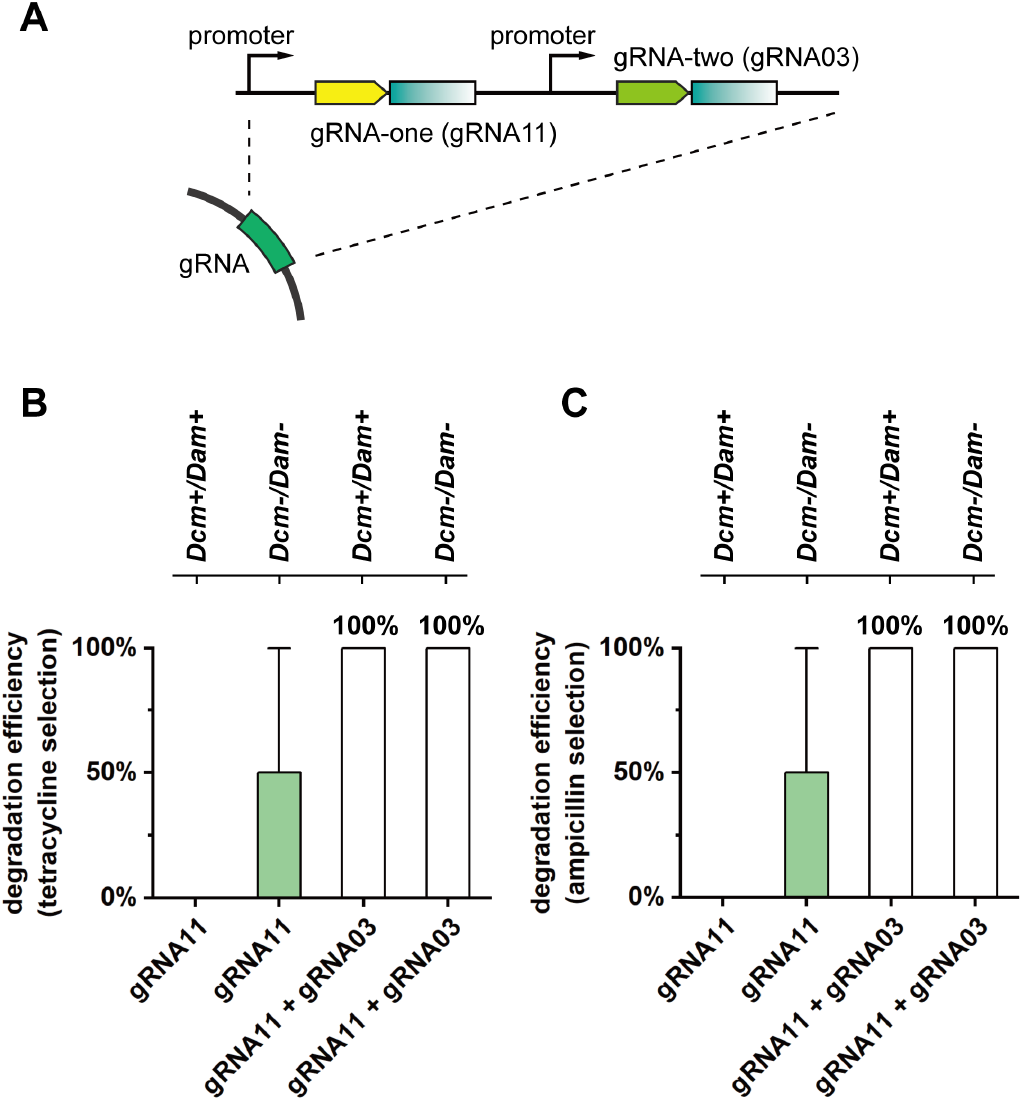
(A) Design scheme of tandem gRNAs for duplex VADER system with gRNA11 and gRNAO3 as an example. Degradation efficiencies of VADER plasmids pEL11 (with gRNA11) and pELdu (with tandem gRNA11 and gRNA03) for *tetA* degradation (B) and plasmid elimination (C) in *E. coli* with functional methylation mechanisms and with methylation deficiency.

### Integrating IncP conjugation module for delivery

Transformation is, however, not suitable for the delivery of VADER in the real world due to the requirement of cell competence. One alternative is transduction depending on virus-mediated delivery, for instance, with bacteriophages (22). Though transduction showed great potential in medical applications and eukaryotic gene-editing, it is often highly host-specific and has a limitation in cargo sizes. Conjugation is another possible way for delivery of genetic systems, as it has a broad spectrum of recipient cells, and the feasibility has been demonstrated in principle (22, 23, 34). By applying conjugation machinery, when the donor cell with VADER contacts with the recipient cell harboring ARG, it delivers VADER and then VADER will degrade the target ARG in vivo **(Figure 4A)**.

**Figure 4.**
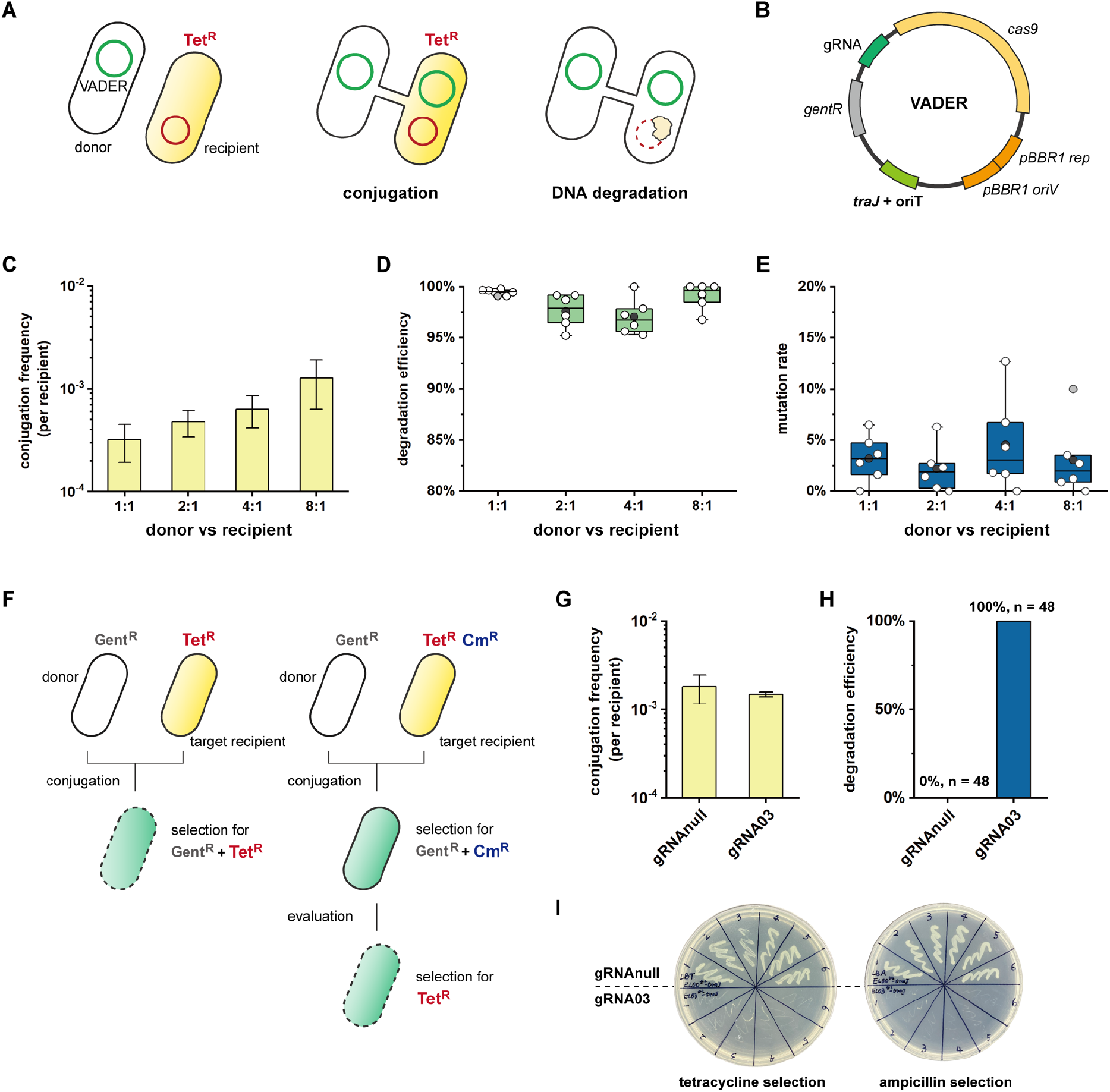
Working mechanism of conjugative delivery of VADER (A) and rational design of artificial IncP modules *(traJ* and *oriT)* (B). (C) Conjugation frequency of VADER evaluated by a non-targeting VADER plasmid pELnc. (D) Degradation efficiency of VADER plasmid pEL03c at different ratio between donor and recipient cells. (E) Mutation rate of a non-targeting VADER plasmid pELnc determined by morphology of colonies and double checked by PCR and sequencing. (F) Experimental scheme for validation of VADER. For a direct selection by gentamycin (Gent^R^) and tetracycline resistances (Tet^R^), the transconjugants receiving a targeting VADER will, by design, be dead which making direct evaluation of VADER impossible. With the assistance of an additional plasmid harboring a third selection criterion (Cm^R^), the transconjugants with a targeting VADER can be selected first by Gent^R^ and Cm^R^, then being evaluated on a second round by tetracycline resistance (Tet^R^). (G) Conjugation frequency of a targeting VADER evaluated directly by selection by Gent^R^ and Cm^R^. (H) Degradation efficiency of VADER evaluated by a second round of selection on Tet^R^. Forty-eight colonies were randomly picked for evaluation (n = 48). (I) Demonstration of VADER on selection plates.

The efficiency of VADER cannot be directly calculated, as all the cells receiving a targeting VADER, as designed, will be dead when being selected by the target ARG **(Figure 4F)**. Therefore, the degradation efficiencies were calculated by comparing targeting and non-targeting VADER, based on the assumption that targeting and non-targeting VADER plasmids have similar conjugation frequencies. For a more concrete and direct validation, we transformed an assistant plasmid pCmR with chloramphenicol resistance (Cm^R^) for the selection of living cells receiving a targeting VADER before testing for the target ARG. With this experimental design, we were able to select cells with Cm^R^, harbored by the target cells besides the target ARG, and Gent^R^, harbored by VADER plasmids, to get colonies receiving VADER plasmids. Then, the colonies were tested for Tet^R^ to evaluate the function of VADER for ARG degradation **(Figure 4F)**. The results showed that the targeting and non-targeting VADER showed similar conjugation frequency **(Figure 4G)**, which confirmed our assumption and data, and the targeting VADER exhibited high degradation efficiency **(Figure 4H and 4I)**. Notably, we observed 100% efficiency in *tetA* degradation and did not find escaper cells from the targeting VADER, and no insertions in the non-targeting VADER were identified. Taking together, VADER can efficiently degrade plasmid-borne ARGs via conjugative delivery.

### A prototype conjugation reactor for VADER

Environmental processes, especially wastewater treatment processes, contain different sectors and remediate contamination or recycle resources in designed units. For instance, the anaerobic-anoxic-aerobic system shows process-level advantages to simultaneously remove organic contaminants, nitrogen and phosphorus by setting up anaerobic, micro-aerobic, and aerobic regions (35). At the process level, it is rational and possible to expand conventional processes by instrumenting a specialized sector for ARG degradation, and we propose a conjugation reactor as the additional unit to biological processes. The conjugation reactor contains donor cells with the targeting VADER, and it receives microbes from the biological processes. When these microbes enter the reactor, VADER will be delivered to degrade the target ARGs. Finally, the ARG-free “clean” microbes can return to their assigned bioprocess, and it will be safer for final disposal when required (e.g., sludge reduction and landfilling).

As a proof of principle, we built a milliliter-level conjugation reactor **(Figure 5A)**, and the conjugation reactor works in a sequencing batch mode. With pre-cultivated donor cells inside the reactor, the process can be divided into five stages: **1)** the target recipient cells as inlet enter the reactor, **2)** stirring to mix the donor and recipient cells, **3)** conjugation takes place in the reactor, **4)** vigorous stirring to separate conjugated cells, and **5)** discharge of ARG-free cells **(Figure 5B)**. We tested our design using *E. coli* S17-1 with VADER plasmid pEL03c as the donor and *E. coli* with pBR322 and pCmR as the target recipient. pCmR was included to enable the evaluation of ARG degradation with living cells **(Figure 4F)**. By setting 8:1 as the ratio between the donor and recipient, we found efficient delivery of VADER (1.93 × 10^−4^ transconjugants per recipient) after 2 h of conjugation, and the conjugation frequency reached 3.14 ×10^−3^ transconjugants per recipient at 3 h **(Figure 5C)**. No longer than 3 h of evaluation was performed to minimize the exaggeration effect caused by cell growth. Then, we randomly picked colonies of recipients with the working VADER after 2 h and 3 h of conjugation, respectively, and tested for the Tet^R^. As expected, all tested cells cannot grow on the selection plates, indicating successful degradation of the target ARG with 100% efficiencies **(Figure 5D)**.

**Figure 5.**
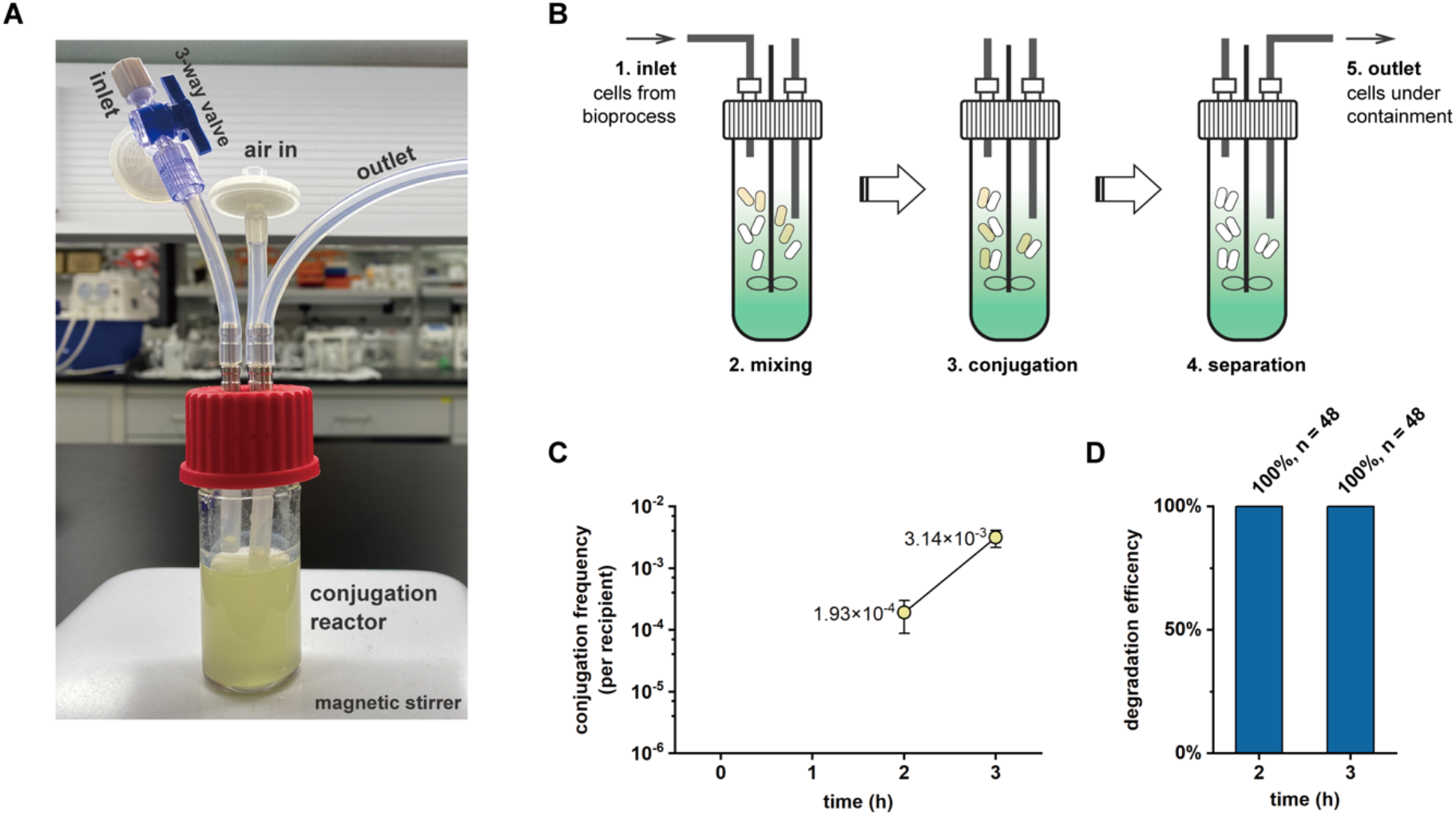
(A) Configuration of the prototype conjugation reactor with a total volume of 30 mL and a working volume of 13.5 mL. (B) Operation scheme of the conjugation reactor follows a sequencing batch mode in five stages: inlet, mixing, conjugation, separation, and outlet. (C) Conjugation frequency of VADER performed in the conjugation reactor. Samples were collected at different time points and cultivated in the non-selective medium for one hour before plating. (D) Degradation efficiency of VADER in the reactor. The efficiency was calculated by testing the tetracycline resistance of randomly picked colonies (n = 48) of transconjugants with VADER.

## Discussion

CRISPR-Cas-based living therapeutics have been thriving in enterprising medical research and applications (36, 37). Different from medical applications where CRISPR-Cas systems can be carried by, i.e., probiotics, for the administration in patients, environmental biotechnology takes place in reactors, lagoons, or other civil facilities, making it quite different and challenging to find practical scenarios for genetic systems like CRISPR. The challenges lay in **1)** designing an easy-to-operate and simple system, **2)** scaling up the delivery, and **3)** inhabiting the system at the process level. As such, we demonstrated here in principle that the combination of tailored CRISPR-Cas immunity and IncP machinery can reside in a conjugation reactor for ARG degradation with environmental purposes.

First, we developed VADER as a one-plasmid system with constitutive functions, leading to the inactivation of target ARG and elimination of the carrying plasmids in a single step. To overcome the toxicity of the constitutive CRISPR-Cas system in bacteria, we optimized the promoter driving gRNAs by mutating the TATA box for the generation of working VADERs. Then, the conjugative delivery of VADER was realized not only on a microliter scale, as commonly being described in labs or in medical applications (23), but also in a reactor, where the conjugation procedures can be sequenced as operational steps. Although our prototype reactor only has a working volume of 13.5 mL, which is way smaller than a real-world reactor, we scaled up the conjugation system from a microliter scale to a 10-mL-scale by over 100 times. While this is a small step toward instigating a practical conjugation reactor in environmental processes, for the first time, we put the CRISPR-Cas system itself rather than a CRISPR-engineered organism in a reactor with operationally conjugative delivery at the process level, unveiling the great potential of the implementation of synthetic biology in environmental processes.

Meanwhile, we do not want to overemphasize our work or omit the drawbacks of VADER. We admit that: **1)** the conjugation frequency is still not high enough for practical applications, **2**) mutations in VADER plasmids during conjugation can still occur, and **3)** the conjugation reactor is currently a prototype under sub-optimal conditions. Further work will be taken place in developing more efficient and stable conjugation systems by, for instance, reprogramming the conjugation machinery, and in optimizing and scaling up the conjugation reactor. Another concern is the generalization of VADER, as a great variety of ARGs may exist in the environment (4, 6, 8) whereas CRISPR-Cas immunity is specialized in precise targeting. Since the CRISPR-Cas immunity works in a sequence-driven manner regardless of the forms and types of DNAs, the utilization of VADER for diverse ARGs may be achievable by targeting conserved DNA sequences in different ARGs or in the plasmids. But still, a foreseen practical demonstration may be deploying our design in the pharmaceutical industry with relatively defined types of antibiotics and ARGs, where the environmental contradiction is also more appealing. Finally, VADER is a genetic system with a selection marker and a foreign CRISPR-Cas system as the functional module. So, VADER per se requires a fine control to prevent leakage and distribution by combining physical containment of the reactor and pre-installed biocontainment modules (38, 39).

With the rising of synthetic biology, genetic materials, including but not limited to ARGs, have been used for generating genetically modified organisms and artificial genetic systems in all aspects of human lives (38, 40), committing a high probability of spread in the environment. Genetic materials, especially DNAs, are biomolecules and more, where the DNA sequences determine their functions and impacts (41). They are stable, inheritable, transferable, and sometimes self-recoverable, but the environmental and ecological impacts are still mysteries (41, 42). While pre-installed or external biocontainment approaches have been investigated and instigated to warrant biosafety, environmental processes, as key barriers, will inevitably be challenged by undesired natural and synthetic genetic materials in the near future. Therefore, a new sector in the framework of environmental processes specialized in the removal of genetic materials or genetic pollutants is crucial, and our work gives a paradigm for this endeavor. On the path towards a healthy and sustainable future, we believe that the cross-disciplinary advantages of synthetic biology and environmental biotechnology are just awakening.

## Materials and Methods

### Stains and Media

*Escherichia coli* MG1655 (CGSC#6300) was used as a host of plasmid-borne ARG for evaluating VADER with functional methylation mechanisms. *E. coli* HST04 (Takara Bio.) was used as the methylation deficient strain to assess the effects of methylation on CRISPR-Cas mediated ARG degradation. *E. coli* S17-1 harboring the complete IncP conjugation machinery in the chromosome was used as the donor strain for conjugation (22, 34). *E. coli* DH5α (Takara Bio.) was used for molecular cloning to generate and maintain plasmids. *E. coli* strains were cultivated in Luria-Bertani (LB) medium with ampicillin (100 μg/L), gentamycin (20 μg/L), tetracycline (25 μg/L), and chloramphenicol (34 μg/L) when necessary.

### Plasmid Construction

The plasmids used in this study are summarized in Table S1. pBBR1MCS5 was a lab stock plasmid (25), and pCas9 (Addgene#42876) was a gift from Luciano Marraffini (43). pMV-gRNA with gRNA cassette was synthesized by Beijing Liuhe BGI, and pBR322 was a commercial plasmid from Takara Bio. The first gRNA plasmid pgRNA01 carrying gRNA01 cassette was generated via inverse PCR. pCasEnv was constructed via DNA assembly with pBBR1MCS5 as the backbone and *cas9* from pCas9, and pBBR-LX01 was assembled with pBBR1MCS5 as the backbone and gRNA01 cassette. The first VADER plasmid harboring *cas9* and gRNA01 cassette was generated by fusing gRNA01 cassette from pgRNA01 to pCasEnv. Then, all VADER plasmids were directly constructed via inverse PCR to replace spacers. pELdu was constructed by fusing gRNA03 cassette to pEL11. pELnc and pEL03c were built by fusing *traJ* and *oriT* modules to pELnu and pEL03, respectively. All DNA assemblies used in plasmid generation were following protocols from the merchandiser (Takara Bio.), and inverse PCRs were performed as normal PCRs but with back-to-back primer pairs. Plasmid extraction and purification were conducted using commercial kits from Tiangen Biotech. PCRs were performed with PrimeSTAR Max DNA Polymerase from Takara Bio. All primers used in this study were synthesized by Beijing Liuhe BGI and are listed in Table S2. The sequences of synthesized fragments are listed in Table S3. The gRNAs with spacers and PAMs are summarized in Table S4.

### Bacterial transformation

The preparation of chemically competent cells and *E. coli* transformation were following the standard methods with minor modifications. For preparing competent cells, overnight culture of *E. coli* was inoculated to LB medium and grew until OD_600_ reaching 0.3. The cells were harvested at 4 °C and resuspended in 0.1 M of ice-cold CaCl_2_, staying on ice for 30 min. After repeating the step, cells were resuspended and aliquoted in 0.1 M of CaCl_2_ with 15% (v/v) of glycerol for storage at −80 °C. For transformation, the competent cells were first thawed on ice, and DNA was then added to the cells, followed by incubation on ice for 30 min and a 60 s of heat shock at 42 °C. After recovery in LB medium for 1 h, all cells were centrifuged and resuspended in 100 μL of sterile water before plating on selection plates.

### Bacterial conjugation

*E. coli* conjugation was performed in a 1.5-mL Eppendorf tube at a volume of 100 μL. First, donor and recipient cells were inoculated from overnight cultures and cultivated until OD_600_ reaching 0.8 – 1.0. Then, the donor and recipient cells were washed with LB medium, mixed together, and kept at 37 °C for conjugation. After 1.5 h of conjugation, fresh LB medium was added and cultivated for another 1.5 h of recovery, then plating on selection plates with appropriate antibiotics. Colony-forming units were measured to determine the numbers of transconjugants and recipients. Conjugation frequency was calculated according to the following equation *(eq.1)*.

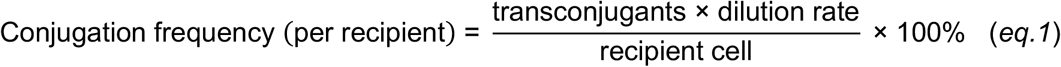

### Analysis for degradation efficiency

Degradation efficiency was evaluated by checking transformants with VADER plasmids on selection plates with the corresponding antibiotic of the target ARG. In the present study, transformants were first selected on gentamycin plates to ensure the transformation of VADER plasmids. Then, at least 12 colonies from each of the plates were randomly picked and re-streaked on plates with tetracycline and ampicillin, respectively, to evaluate the degradation of *tetA* and the integrity of target plasmids. Biological duplication or triplication was performed. The degradation efficiency was calculated with *eq.2*.

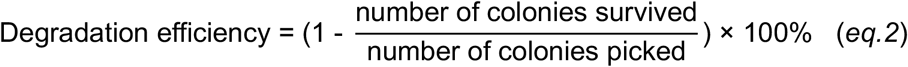

For conjugation experiments, the transconjugants were selected on double selective plates with gentamycin and tetracycline. As designed, transconjugants receiving a targeting VADER, with gRNA targeting *tetA*, will be dead and only escaper cells will survive. Thus, the degradation efficiency for conjugation experiments directly evaluated with tetracycline was calculated by comparing the conjugation frequencies between targeting and non-targeting VADER plasmids based on the hypothesis that they have similar conjugation frequencies.

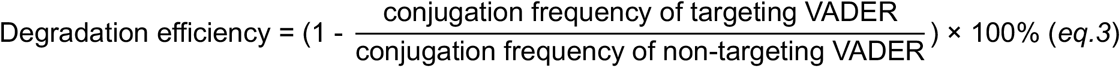

With the assistance of pCmR giving an extra selection criterion, the transconjugants were able to be selected with gentamycin and chloramphenicol. The transconjugants were then tested with tetracycline to evaluate the VADER system. Similarly, a minimum of 12 colonies from the plates with gentamycin and chloramphenicol after conjugation were randomly picked and re-streaked on plates with tetracycline, and the number of survival colonies were recorded for calculation with *eq.2*.

### Colony PCR

A colony was picked, resuspended in sterile water and then inoculated at 100 °C for 10 min. One microliter of the supernatant was used as the template for PCR amplification of designed fragments on pBR322. All primers used for colony PCR are listed in Table S2.

### Prototype conjugation reactor

The conjugation reactor was demonstrated in a 30-mL glass bottle with inlet, outlet, and ventilation tube for air exchange. The reactor was operated as a sequencing batch reactor at 37 °C with a ratio of 8:1 between donor and recipient cells, including five steps: inlet, mixing, conjugation, separation, and outlet. Samples were taken at time points 0, 1 h, 2 h, and 3 h, and the samples were cultivated with fresh LB medium for one hour of recovery before plating to get survival colonies on selection plates. Then, the colonies were tested for tetracycline resistance to evaluate the degradation of *tetA*.

## Supplementary materials

Summary of plasmids and primers used in this study; summary of synthesized DNA fragments; summary of gRNAs with protospacer and PAMs; figures supporting the main results.

## Acknowledgments

This work was supported by the National Natural Science Foundation of China (U20A20146), the Distinguished Young Scholar Program of Shandong Province (Overseas) (2022HWYQ-017), and the Qilu Young Scholar Program of Shandong University (to P.-F.X.).

