## Supplementary materials for "Tailoring CRISPR-Cas Immunity for the Degradation of Antibiotic Resistance Genes"

**Table S1.** Plasmids used in this study.

| **Name** | **Description** | **References** |
|---|---|---|
| pBBR1MCS5 | *pBBR* ori, *gentR* | Lab stock |
| pMV-gRNA | *pUC* ori, gRNA, *bla* | Beijing Liuhe BGI. |
| pBR322 | *colE1* ori, *bla*, *tetA* | Takara Bio. |
| pCas9 | *p15a* ori, *cas9, cmR* | W. Jiang et al. (1) |
| pgRNA01 | pMV-gRNA, gRNA01 | This study |
| pBR322-pUC | pBR322, *pUC* ori | This study |
| pBR322-p15A | pBR322, *p15A* ori | This study |
| pCasEnv | pBBR1MCS5, *cas9* | This study |
| pBBR-LX01 | pBBR1MCS5, gRNA01 | This study |
| pELnu | pCasEnv, gRNAnull | This study |
| pEL01 | pCasEnv, gRNA01 | This study |
| pEL03 | pCasEnv, gRNA03 | This study |
| pEL06 | pCasEnv, gRNA06 | This study |
| pEL11 | pCasEnv, gRNA11 | This study |
| pEL12 | pCasEnv, gRNA12 | This study |
| pEL13 | pCasEnv, gRNA13 | This study |
| pEL14 | pCasEnv, gRNA14 | This study |
| pELdu | pCasEnv, gRNA11, gRNA03 | This study |
| pELnc | pCasEnv, gRNAnull, *traJ, oriT* | This study |
| pEL03c | pCasEnv, gRNA03, *traJ, oriT* | This study |
| pCmR | pCas9, *Δcas9* | This study |

**Table S2.** Primers used in this study.

| Primer | Sequence |
|---|---|
| **Primers for In-Fusion DNA assembly** | |
| XIA-LX-001 | TGAGTTAGCTCACTCATTAGGGTCGACGCTGTTTTGAATGGTTCCAA |
| XIA-LX-002 | CGCTTACAATTTCCATTCGCTACTTCAGTCACCTCCTAGC |
| XIA-LX-003 | GCTAGGAGGTGACTGAAGTAGCGAATGGAAATTGTAAGCG |
| XIA-LX-004 | TTGGAACCATTCAAAACAGCGTCGACCCTAATGAGTGAGCTAACTCA |
| XIA-LX-023 | GAGTTAGCTCACTCATTAGGTGTCTTGCGTCTCTGTCGAC |
| XIA-LX-024 | CCATTCAAAACAGCGTCGACTCTGGTACATGCACCGACTC |
| XIA-LX-025 | GAGTCGGTGCATGTACCAGAGTCGACGCTGTTTTGAATGG |
| XIA-LX-026 | GTCGACAGAGACGCAAGACACCTAATGAGTGAGCTAACTC |
| XIA-LX-015 | ATGTACCAGATCGCTAACTGGCGAATGGAAATTGTAAGCG |
| XIA-LX-016 | CAGTTAGCGATCTGGTACATCCATTCAAAACAGCGTCGAC |
| XIA-LX-030 | GAACATGTGAGCAAAAGGCC |
| XIA-LX-031 | AGGATCTAGGTGAAGATCCT |
| XIA-LX-032 | AGGATCTTCACCTAGATCCT |
| XIA-LX-033 | GGCCTTTTGCTCACATGTTC |
| XIA-LX-034 | GAACATGTGAGCAAAAGGCCTGGAAGATGCCAGGAAGATA |
| XIA-LX-035 | AGGATCTAGGTGAAGATCCTCTGACTTCAGGTGCTACATT |
| XIA-LX-099 | CGGTGCATGTACCAGAGTCGACTTGACAGCTAGCTCAGTC |
| XIA-LX-100 | TTGGAACCATTCAAAACAGCGCACCGACTCGGTGCCAC |
| XIA-LX-101 | GACTGAGCTAGCTGTCAAGTCGACTCTGGTACATGCACCG |
| XIA-LX-102 | GTGGCACCGAGTCGGTGCGCTGTTTTGAATGGTTCCAA |
| XIA-LX-081 | ACCGAACAACTCCGCTTCGGGGTCATTATA |
| XIA-LX-082 | AGATCGGCTTCCCGGCCTCTTCTTGATGGAGCGCAT |
| **Primers for inverse PCR** | |
| XIA-LX-027 | ATGTACCAGATCGCTAACTGGTTTTAGAGCTAGAAATAGC |
| XIA-LX-028 | CAGTTAGCGATCTGGTACATGCTAGCATTATACCTAGGAC |
| XIA-LX-009 | TTGAAGGCTCTCAAGGGCATGTTTTAGAGCTAGAAATAGC |
| XIA-LX-010 | ATGCCCTTGAGAGCCTTCAAGCTAGCATTATACCTAGGAC |
| XIA-LX-036 | TGGCGACCACACCCGTCCTGGTTTTAGAGCTAGAAATAGC |
| XIA-LX-037 | CAGGACGGGTGTGGTCGCCAGCTAGCATTATACCTAGGAC |
| XIA-LX-073 | CATCCAGGGTGACGGTGCCGGTTTTAGAGCTAGAAATAGC |
| XIA-LX-074 | CGGCACCGTCACCCTGGATGGCTAGCATTTACCTAGGAC |

| | |
|---|---|
| XIA-LX-083 | ATGCCCTTGAGAGCCTTCAAGCTAGCATTAACCTAGGAC |
| XIA-LX-084 | ATGCCCTTGAGAGCCTTCAAGCTAGCATTTACCTAGGAC |
| XIA-LX-069 | CATGATCGCGTAGTCGATAGGTTTTAGAGCTAGAAATAGC |
| XIA-LX-096 | CTATCGACTACGCGATCATGGCTAGCATTTACCTAGGAC |
| XIA-LX-071 | ACCGTCACCCTGGATGCTGTGTTTTAGAGCTAGAAATAGC |
| XIA-LX-097 | ACAGCATCCAGGGTGACGGTGCTAGCATTTACCTAGGAC |
| XIA-LX-067 | GAAGATCGGGCTCGCCACTTGTTTTAGAGCTAGAAATAGC |
| XIA-LX-086 | AAGTGGCGAGCCCGATCTTCGCTAGCATTTACCTAGGAC |
| XIA-LX-128 | CAGTTAGCGATCTGGTACATGCTAGCATTTACCTAGGAC |
| XIA-LX-136 | ATGTCATGATAATAATGGTTGATACTTCTATTCTACTCTGACTGC |
| XIA-LX-137 | CCATTATTATCATGACATGATGACGACCATCAGGGACA |
| **Primers for colony PCR and sequencing** | |
| XIA-LX-029 | TATGCATGCGCCCAATAC |
| XIA-LX-089 | ATATCGGCACAAATAGCGTC |
| XIA-LX-090 | TTTGTCATTGGGTTTGACCC |
| XIA-LX-091 | GCATGGATGACTCGGAAGTC |
| XIA-LX-092 | ACAAGTGTCTGGACAAGGCG |
| XIA-LX-093 | GGCACAAATTTTGGATAGTC |
| XIA-LX-094 | CCGTTAAAGAGTTACTAGGG |
| XIA-LX-144 | GGCATAGGCTTGGTTATGCC |
| XIA-LX-145 | ATCTTCCCCATCGGTGATGT |
| XIA-LX-146 | CGGGGAGGCAGACAAGGTAT |
| XIA-LX-147 | CGCGGTATTATCCCGTGTTG |
| XIA-LX-148 | GTCACGCTCGTCGTTTGGTA |

**Table S3.** Sequences of synthesized genetic modules.

| Gene | Sequence [a] |
|---|---|
| gRNA cassette | GTCGACGGATCCTTGACAGCTAGCTCAGTCCTAGGTATAATGCTAGCA<br>ATAGTCGGTGATTTCTTCGGTTTTAGAGCTAGAAATAGCAAGTTAAAAT<br>AAGGCTAGTCCGTTATCAACTTGAAAAAGTGGCACCGAGTCGGTGCAT<br>GTACCAGATCGCTAACTGGTTGCAAATAAAACGAAAGGCTCAGTCGAA<br>AGACTGGGCCTTTCGTTTTATCTGTTGTTTGTCGGTGAACGCTCTCGT<br>CGAC |
| *traJ* and *oriT* [b] | TTCGGGGTCATTATAGCGATTTTTTCGGTATATCCATCCTTTTTCGCAC<br>GATATACAGGATTTTGCCAAAGGGTTCGTGTAGACTTTCCTTGGTGTAT<br>CCAACGGCGTCAGCCGGGCAGGATAGGTGAAGTAGGCCCACCCGCG<br>AGCGGGTGTTCCTTCTTCACTGTCCCTTATTCGCACCTGGCGGTGCTC<br>AACGGGAATCCTGCTCTGCGAGGCTGGCCGGCTACCGCCGGCGTAA<br>CAGATGAGGGCAAGCGGATGGCTGATGAAACCAAGCCAACCAGGAAG<br>GGCAGCCCACCTATCAAGGTGTACTGCCTTCCAGACGAACGAAGAGC<br>GATTGAGGAAAAGGCGGCGGCGGCCGGCATGAGCCTGTCGGCCTAC<br>CTGCTGGCCGTCGGCCAGGGCTACAAAATCACGGGCGTCGTGGACTA<br>TGAGCACGTCCGCGAGCTGGCCCGCATCAATGGCGACCTGGGCCGC<br>CTGGGCGGCCTGCTGAAACTCTGGCTCACCGACGACCCGCGCACGG<br>CGCGGTTCGGTGATGCCACGATCCTCGCCCTGCTGGCGAAGATCGAA<br>GAGAAGCAGGACGAGCTTGGCAAGGTCATGATGGGCGTGGTCCGCC<br>CGAGGGCAGAGCCATGACTTTTTAGCCGCTAAAACGGCCGGGGGGT<br>GCGCGTGATTGCCAAGCACGTCCCCATGCGCTCCATCAAGAAGA |

[a] coding sequences in the synthesized genes are shown in blue. Both synthesized genes were carried by pMV plasmid with $Amp^R$.

[b] *traJ* and *oriT* were synthesized together in one plasmid.

**Table S4.** gRNA sequences used in this study.

| gRNA | Target | Strand [a] | PAM | Protospacer |
|---|---|---|---|---|
| gRNAnull | - | - | - | ATGTACCAGATCGCTAACTG |
| gRNA01 | *tetA* | C | CGG | GCAGCGCTCTGGGTCATTTT |
| gRNA03 | *tetA* | N | CGG | TTGAAGGCTCTCAAGGGCAT |
| gRNA06 | *tetA* | C | TGG | TGGCGACCACACCCGTCCTG |
| gRNA11 | *tetA* | C | CGG | GAAGATCGGGCTCGCCACTT |
| gRNA12 | *tetA* | N | TGG | CATGATCGCGTAGTCGATAG |
| gRNA13 | *tetA* | C | AGG | ACCGTCACCCTGGATGCTGT |
| gRNA14 | *tetA* | N | AGG | CATCCAGGGTGACGGTGCCG |

[a] C stands for coding strand, and N stands for non-coding strand.

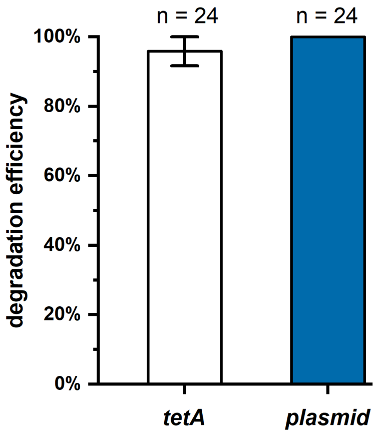


**Figure S1.** Degradation efficiency of pEL14 with mutated J23119 promoter leading the gRNA.

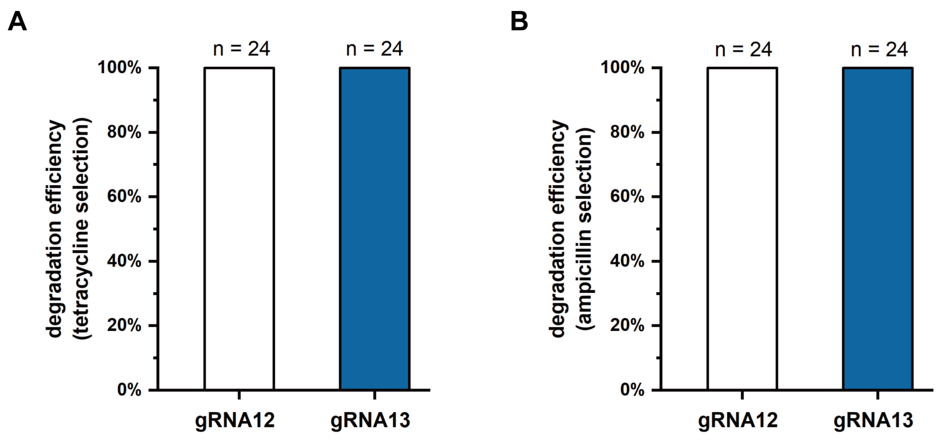


**Figure S2.** Degradation efficiency of pEL12 (with gRNA12) and pEL13 (with gRNA13) with J23119** promoter leading the gRNA cassette. (A) shows the degradation efficiency of *tetA*, and (B) shows the degradation efficiency of plasmid pBR322 by selection on plates with ampicillin.

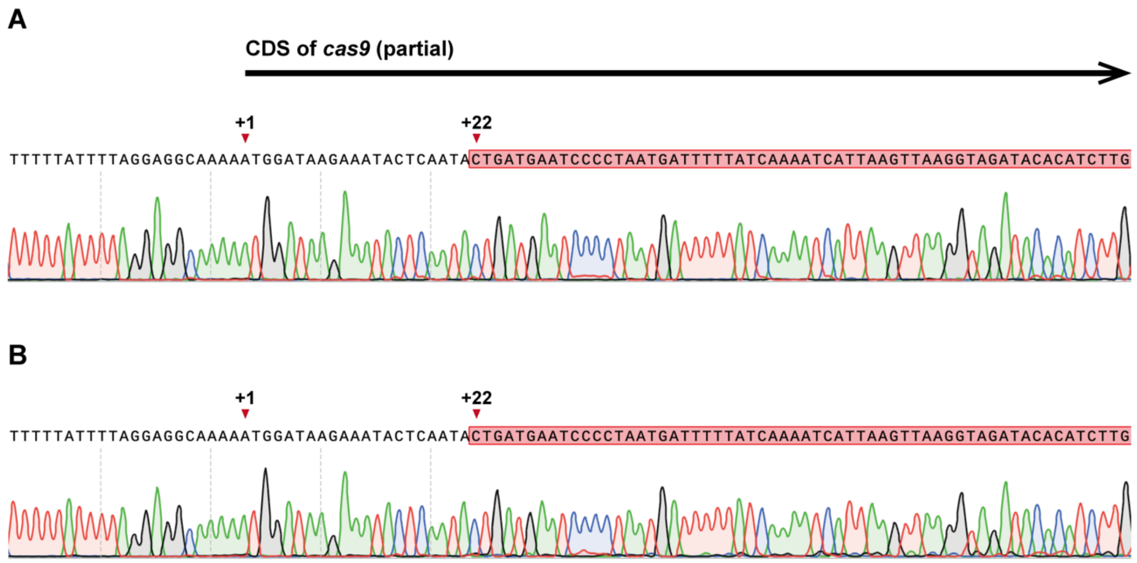


**Figure S3.** Sequencing results of mutated VADER plasmids from escaper cells with pEL03c (A) and from transconjugants from pELnc (B). Position +1 shows the start of the CDS of *cas9*, and the highlighted sequences indicate the inserted IS-4 transposon sequence (partial).
